# Hybrid assembly of the genome of the entomopathogenic nematode *Steinernema carpocapsae* identifies the X-chromosome

**DOI:** 10.1101/571265

**Authors:** Lorrayne Serra, Marissa Macchietto, Aide Macias-Muñoz, Cassandra Joan McGill, Isaryhia Maya Rodriguez, Bryan Rodriguez, Rabi Murad, Ali Mortazavi

## Abstract

Entomopathogenic nematodes from the genus *Steinernema* are lethal insect parasites that quickly kill their insect hosts with the help of their symbiotic bacteria. *Steinernema carpocapsae* is one of the most studied entomopathogens due to its broad lethality to diverse insect species and its effective commercial use as a biological control agent for insect pests, as well as a genetic model for studying parasitism, pathogenesis, and symbiosis. In this study, we used long-reads from the Pacific Biosciences platform and BioNano Genomics Irys system to assemble the best genome of *S. carpocapsae* ALL strain to date, comprising 84.5 Mb in 16 scaffolds, with an N50 of 7.36Mb. The largest scaffold, with 20.9Mb, was identified as chromosome X based on sex-specific genome sequencing. The high level of contiguity allowed us to characterize gene density, repeat content, and GC content. RNA-seq data from 17 developmental stages, spanning from embryo to adult, were used to predict 30,957 gene models. Using this new genome, we performed a macrosyntenic analysis to *Caenorhabditis elegans* and *Pristionchus pacificus* and found *S. carpocapsae’s* chromosome X to be primarily orthologous to *C. elegans’* and *P. pacificus’* chromosome II and IV. We also investigated the expansion of protein families and gene expression differences between male and female stage nematodes. This new genome and more accurate set of annotations provide a foundation for new comparative genomic and gene expression studies within the *Steinernema* clade and across the Nematoda phylum.

**Article Summary:** The insect killing worms Steinernema carpocapsae is a model organism for parasitism and symbiosis. The authors have used long reads and optical mapping to generate substantially contiguous assembly and a new set of gene annotations. They have identified the X chromosome as well as expansions in specific family proteases found in the venom of this worm. A macrosyntenic analysis with *C. elegans* shows a broad conservation of ancestral chromosomes with the exception of chromosome X. This new assembly will be useful to the *Steinernema* community and the broader nematode genomics community.

## Introduction

Nematodes are round, non-segmented worms with a simple body plan that populate all known biological niches. *Caenorhabditis elegans* is by far the best-characterized nematode species and was the first metazoan to have its genome assembled (*C. elegans* Sequencing Consortium, 1998). Studies in *C. elegans* have been crucial for the study of a multitude of biological processes such as development, cell specification, cell differentiation, apoptosis and genome evolution (Deppe et al., 1978; Kaletta and Hengartner 2006; Rothman et al., 2014; Segref et al. 2010). Other free-living, androdiecious nematode genomes such as *Pristionchus Pacificus* and *Panagrellus redivivus* have provided tools for genomic comparisons in developmental processes and phenotypic plasticity (Rödelsperger et al., 2017; Srinivasan et al., 2013).

However, free-living nematodes represent only a subset of the entire nematode phylum as they exclude parasitic nematodes, which are of most agricultural and biomedical interest. Parasitic nematodes have a diverse group of hosts including insects, plants, and humans. *Steinernema carpocapsae* is an Entomopathogenic nematode (EPN) that is a lethal parasite of insects but is not harmful to humans or plants. *S. carpocapsae* is of great interest because of its wide host range with approximately 200 possible insect hosts and the possible orthology of its toxins to mammalian-parasitic nematodes (Shapiro-Ilan et al., 2017; Yang et al., 2015). In many countries, *S. carpocapsae* are commercialized for use as insect pest control (Dillman et al., 2012; Dito et al., 2016). *Steinernema* has a symbiotic relationship with pathogenic *Xenorhabdus* bacteria where they collectively infect and kill a host within a few days, reproduce separately within the host, and reassociate when forming infective juveniles that will seek out the next host (Hirao et al., 1999). *Steinernema* and its bacterial symbiont have been extensively studied as genetic models to explore symbiosis and pathogenesis (Martens and Goodrich-Blair 2005; Sicard et al., 2003). EPNs are also an excellent satellite organism to study mammalian parasitism as they are closely related to the *Strongyloididae* nematodes (Blaxter et al., 1998). In particular, *Steinernema’s* mechanism of host seeking by olfactory and other sensory cues likely offers a great model for mammalian-parasitic nematodes (Gang & Hallem, 2016).

Previously published draft genomes done using Illumina sequencing for 5 *Steinernema* species opened the door to study evolutionary traits among these parasitic nematodes and compare them to free-living nematodes. These genomes helped elucidate gene families involved in parasitism that were expanded in *Steinernema*, such as proteases and protease inhibitors, while a comparative analysis to *C. elegans* revealed orthologous non-coding regulatory motifs (Dillman et al., 2015). *S. carpocapsae* and *S. feltiae* were further used to study the extent of expression conservation in orthologous genes across nematode families that were associated with embryonic development (Macchietto et al., 2017). This study found a funnel-shaped model of embryonic development based on high conservation of orthologous genes between *Caenorhabditis* and *Steinernema* (Macchietto et al., 2017). The draft genomes also allowed for further studies in neuropeptide sensory perception in *S. carpocapsae*, and the study of lethal venom proteins in both *S. carpocapsae* and *S. feltiae* (Lu et al., 2017; Morris et al., 2017).

The draft genome assembly of *S. carpocapsae* ALL strain had 1,578 contigs with an estimated genome size of 85.6 Mb, N50 of 300kb and more than 28 thousand predicted genes (Dillman et al., 2015). Another study, also using short-reads assembled *S. carpocapsae* Breton strain into 347 scaffolds with an N50 of 1.24Mb (Rougon-Cardoso et al., 2016). The draft genome status, although helpful, can hinder further studies of protein families’ expansions involved in parasitism. The high fragmentation also deters the analysis of conserved gene sequences, which could play a significantly larger role in the evolution of this phylum. Therefore, a more contiguous assembly of the *S. carpocapsae* genome will contribute to further understanding the mechanisms of *Steinernema* genome evolution. In this study, we reassembled the genome with long-read sequencing from Pacific BioSciences (PacBio) in conjunction with optical mapping using BioNano Irys system, which produces high throughput physical maps (Lam et al., 2012; Rhoads and Au 2015). The newly assembled genome consists of 16 scaffolds with nearly 31 thousand predicted genes and we identified the largest scaffold as chromosome X. We observed clusters of expansion of proteases families and found gene expression differences in catalytic activity between male and female stage nematodes.

## Results

### *S. carpocapsae* largest scaffold identified as chromosome X

We used PacBio sequencing along with BioNano optical mapping to generate a *S. carpocapsae* genome assembly consisting of 17 scaffolds with a N50 of 7.36Mb and GC content of 45.7%. Comparison of our genome to the Breton strain from Rougon-Cardoso et al., 2016 allowed us to merge our smallest contig of 150Kb with scaffold 7 for a total of 16 scaffolds. *S. carpocapsae* is known to have 5 chromosomes, and the largest scaffold of 20.9Mb, could correspond to an assembled chromosome (Supplemental table 1). We performed Illumina sequencing of the DNA of males and females, being careful to collect females that had not mated to identify scaffolds that would encompass the chromosome X in our new assembly. The largest scaffold displayed a characteristic 2-fold difference in coverage between females (orange) and males (blue), which we therefore renamed as Chromosome X (Figure 1A). Scaffold 15, which is much smaller (207kb or 1% of the size of chromosome X) also showed a similar difference in coverage and is likely a small part of the X chromosome. The remaining 1+Mb scaffolds suggest that we have the remaining four chromosomes in 14 pieces at most.

**Figure 1.**
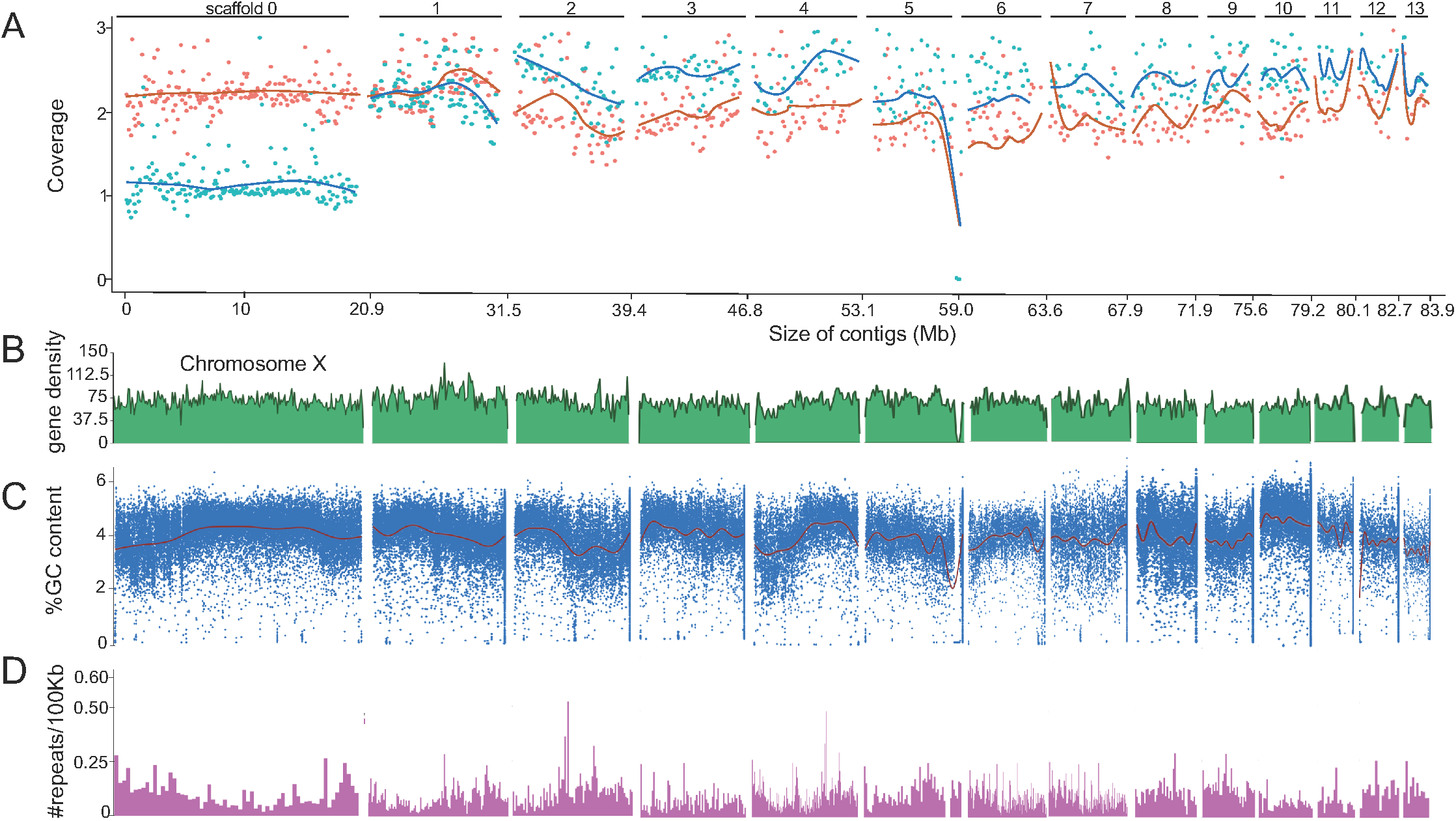
Identifying the X chromosome in *S. carpocapsae*. **(A)** Illumina sequencing of females (orange) and males (blue) shows coverage in order of longest to shortest for scaffolds >1Mb, in which scaffold 0 is chromosome X. **(B)** Gene density calculated for 100kb intervals. **(C)** GC content for every 100kb with a sliding window of 1KB. **(D)** Number of repeats per 100Kb.

We used our RNA-seq transcriptomes from published studies (Marissa Macchietto et al., 2017) and this study to reannotate the genes on this new assembly and identified 30,957 genes using Augustus (see methods). We then calculated the gene density along the scaffolds and observed a uniform distribution of genes on chromosome X, which is similar to observations in *Caenorhabditis elegans* and *Pristionchus Pacificus* (Andersen et al., 2012; Rödelsperger et al., 2017). Gene density varied little along and among all other scaffolds (Figure 1B). Chromosome X has a rich %GC content in its center when compared to its arms. All other scaffolds either have uniform %GC content or show a distinct increase or decrease (Figure 1C). Lastly, we analyzed the repeat content of chromosome X and scaffolds. As expected from other assembled nematode chromosomes, chromosome X repeats are more frequent on the arms than in its central region (Hillier et al., 2007; Rödelsperger et al., 2017; Yin et al., 2018). All other scaffolds that are part of autosomal chromosomes either have uniform repetitive sequences (such as scaffold 3) or scaffolds in which repetitive sequence either decrease or increase (such as scaffolds 2 and 4) (Figure 1D). In summary, we assembled the *S. carpocapsae* genome into 16 scaffolds, 14 of which are greater than 1Mb, including a nearly complete chromosome X that shows similar characteristics to other assembled nematode chromosomes.

### *S. carpocapsae* chromosome X is primarily syntenic to *C. elegans* chromosomes II and IV

We performed a macrosyntenic analysis between *S. carpocapsae* and *C. elegans* genomes by identifying the one-to-one orthologs between both species and plotting these genes according to their position in *S. carpocapsae* scaffolds (Figure 2). We found that *S. carpocapsae* chromosome X is primarily homologous to sections of *C. elegans* chromosomes II, X, IV. *S. carpocapsae* chromosome X has approximately 10Mb from *C. elegans* chromosome II, 7.5Mb from *C. elegans* chromosome IV and 2.5Mb from *C. elegans* chromosome X. The other scaffolds are largely homologous to only one of the *C. elegans* chromosomes. Scaffolds 1, 9, and 11 are homologous to chromosome I. Scaffold 15 is homologous to chromosome II. Scaffolds 4, 7, 8, 13, and 16 are homologous to chromosome III. Scaffolds 2, 6, 10, and 14 are homologous to chromosome IV. Lastly, scaffolds 3, 5, and 12 are homologous to chromosome X.

**Figure 2.**
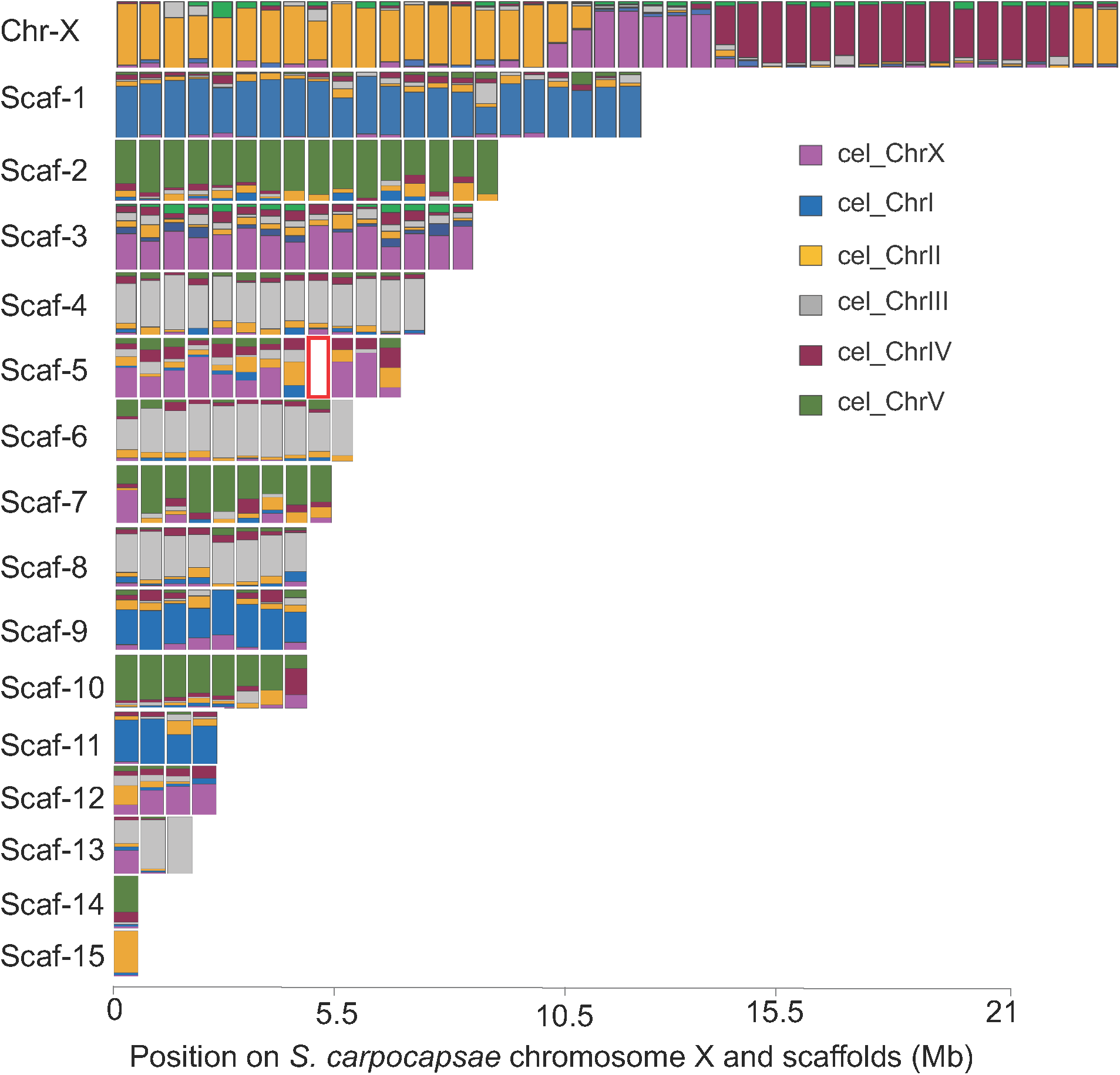
Macrosynteny between *S. carpocapsae* and *C. elegans*. *C. elegans* one-to-one orthologous genes had their position predicted in the *S. carpocapsae* assembly. Each rectangle represents the fraction of *C. elegans* genes present per 500-Kb window in *S. carpocapsae*. Red rectangle indicates no synteny.

Next, we investigated whether *S. carpocapsae* represents the ancestral state by repeating the macrosyntenic analysis with *Pristionchus pacificus* (supplemental Figure 1). The *P. pacificus* chromosome configuration has been reported to represent an ancestral state while *C. elegans* and *Strongylid* nematodes represent the derived state (Rödelsperger et al., 2017). *S. carpocapsae* chromosome X is primarily syntenic to *P. pacificus* chromosomes II, X and IV. The other scaffolds are largely homologs to only one of *P. pacificus* chromosomes. Scaffold 15 is a homolog to chromosome II. Scaffolds 2, 3, 5, 6, 7, 10, 12, and 14 are homologs to chromosome I. Scaffolds 4, 8, and 13 are homologs to chromosome III. Lastly, scaffolds 1, 9 and 11 are homologous to chromosome V. *P. pacificus* chromosomes II, III, IV, X are homologous to the same chromosomes in *C. elegans* (Rödelsperger et al., 2017). However, *P. pacificus* chromosome I is partially *C. elegans* chromosome V and X. *S. carpocapsae* chromosome X configuration is likely similar between *C. elegans* and *P. pacificus* because of the homology between these species for chromosomes II, IV, X. *S. carpocapsae* scaffolds do not share orthologous regions with chromosome X of *P. pacificus.* However, several scaffolds are orthologs with chromosome I (supplemental Figure 2). In summary, *S. carpocapsae* chromosome X is homologous to *C. elegans* and *P. pacificus* chromosomes II, IV, and X. Therefore, *S. carpocapsae* have a derived state and translocation has occurred in the branch leading to the ancestors of *Steinernema*.

### Rapid metalloprotease expansion in S. carpocapsae

A previous study comparing nematode genomes found an expansion of proteases and protease inhibitors in *Steinernema* (Dillman et al., 2015). *S. carpocapsae* in particular had 654 peptidases, approximately one-third of them being metalloproteases and one-third being serine proteases (Dillman et al., 2015). It was suggested that the potential function of these genes may be to aid in parasitism (Dillman et al., 2015). Further studies on the mechanisms of *S. carpocapsae* infection identified 472 venom proteins that included proteases and protease inhibitors (Lu et al., 2017). Our new genome allows us to investigate potential molecular mechanisms by which these gene families may be expanding. In addition, an improved assembly allows us to determine the location of these genes and these genes are in fact numerous individual unique genes or misassembled genes.

In our assembly, we identified 228 unique metalloproteases and 254 serine proteases. Mapping the genes to their chromosome locations reveals aggregation of gene clusters (Figure 3A-B). In order to visualize the location of venom genes, we colored them in red. Venomous metalloproteases form a cluster on scaffold 7, while venomous serine proteins cluster on scaffolds 5, 8 and 14 (Figure 3A-B). The location of these expanded genes suggest that they are evolving by tandem duplications. One-to-one orthologs to *C. elegans* shows that the 228 metalloprotease genes correspond to 90 *C. elegans* gene models and the 284 serine proteases correspond to 50 genes in *C. elegans*. The phylogenetic trees for these gene families inform us about their potential molecular history and sequence similarity. While many of the metalloproteins have orthologs in *C. elegans* with a few duplications, serine proteins seem to have undergone and extensive and rapid expansion in *S. carpocapsae.* Interestingly, there is a small grouping of metalloprotease duplicates that belong to the cluster on chromosome 7; these venomous genes all contain an M14A domain (Figure 2C). Similarly, many of the venomous serine proteases group within what we refer to as the “serine expansion”; these genes have the S01A domain (Figure 3D).

**Figure 3.**
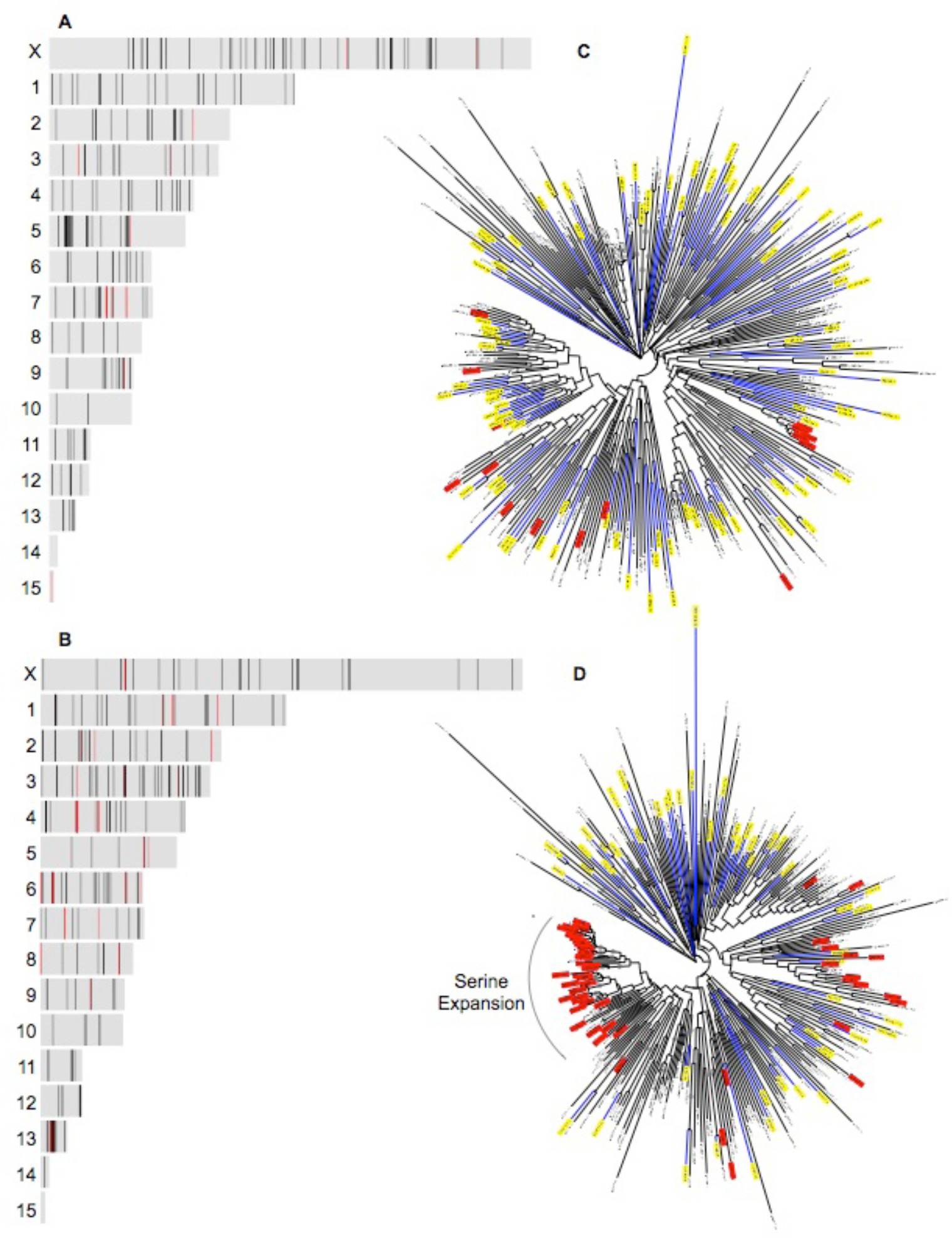
Protease expansion in *S. carpocapsae*. **A)** Location of metalloprotease genes on *S. carpocapsae* scaffolds. Darker bands represent gene overlap or gene clusters. Red bands represent venomous genes. **B)** Location of serine proteases on *S. carpocapsae* scaffolds. Darker bands represent gene overlap or gene clusters. Red bands represent venomous genes. **C)** Maximum-likelihood phylogenetic tree of metalloproteases in *S. carpocapsae* and their orthologs in *C. elegans* (highlighted yellow) using a bootstrap support of 100 replicates. Gene names highlighted in red represent venomous metalloproteases. **D)** Maximum-likelihood phylogenetic tree of serine proteases in *S. carpocapsae* and their orthologs in *C. elegans* (highlighted yellow) using a bootstrap support of 100 replicates. Gene names highlighted in red represent venomous serine protease

### Gene expression profiling *eleven* post-development stages reveals sex specific differences

We next characterized the transcriptome of 11 post-developmental stages with single-nematode RNA-seq analysis in order to identify the similarities between transcriptional profiles of early post-activation, late post-activation, and adults (Serra et al., 2018). Nematodes were activated *in-vitro* on plates with lawns of *X. nematophila*. We collected early post-activation stages nematodes at 3, 6, 9, 12, and 15 hours as well as later post-activation time points that are thought to correspond to later larval stages such as 24, 36, and 48 hours as well as adult females and males. We first generated a heatmap of all genes expressed with a minimum of 1 Transcript Per Million(TPM) (Figure 4A). Early and late post-activation have distinct gene expression profiles, but late post-activation has a similar gene expression profile to female and male adults. Interestingly, we found that two of the nematodes collected at 24 hours have a transcriptional profile similar to adult females, whereas the third nematode collected at the same hour is more similar to an adult male. Similarly, we found that two of the 36 hours nematodes have a male-like profile and one nematode has a female-like profile. At 48 hours all nematodes have a female transcriptome profile, which we assume simply represents our chance of collecting three out of three female samples randomly.

**Figure 4.**
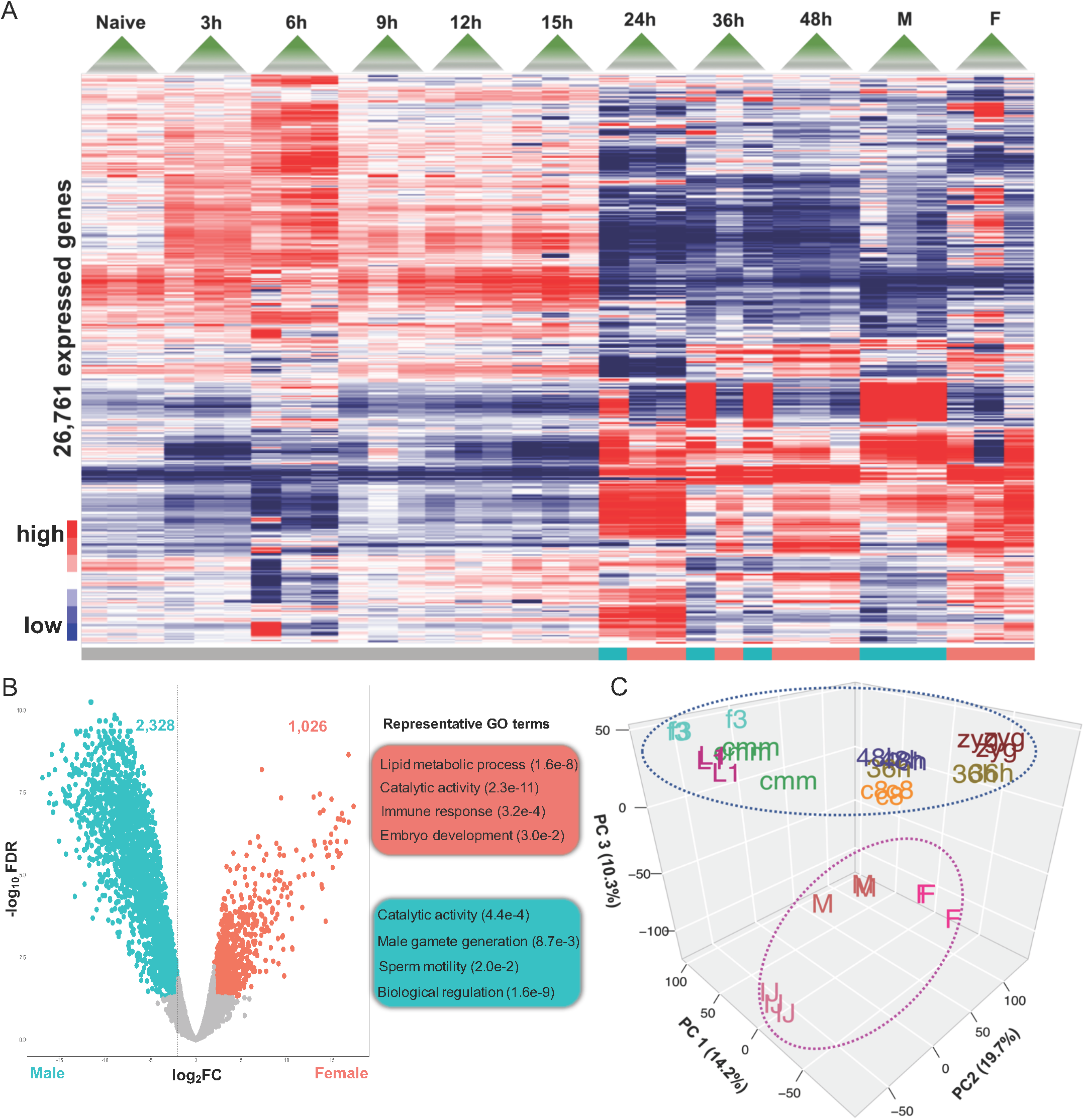
Gene expression profiling of *S. carpocapsae* development in 11 embryonic stages and 11 post-IJs development. **(A)** Heatmap showing 26,761 genes with minimum of 1 transcript per million (TPM). **(B)** Volcano plot of 12,461 differentially expressed genes with minimum of 1TPM, between male (blue) and female (orange) and representative GO terms. **(C)** Principal Component Analysis of all genes expressed in embryos and during an activation time course. Developmental stages are separated from major life stages (IJ, adults) by PC3.

We then conducted a comparative gene expression analysis of adult males and females. We found 2,328 genes downregulated and 1,026 upregulated between males and females (Figure 4B). GO terms for the 2,328 upregulated genes showed strong enrichment for lipid metabolic process (p-value 1.6e-8), catalytic activity (p-value 2.3e-11), immune response (p-value 3.2e-4) and embryo development (p-value 3.0e-2). In contrast, the 1,026 downregulated genes are related to male gamete generation (p-value 8.7e-3), sperm motility (2.0e-2), biological regulation (p-value 1.6e-9) and catalytic activity (p-value 4.4e-4). We generally recovered distinct major GO term categories between female and male except for the category “catalytic activity”, which led us to investigate whether the GO terms for females and males are enriched for distinct enzymes. Male GO terms were enriched for phosphotransferase activity (p-value 5.93e-94), kinase activity (p-value 1.14e-23), phosphatase activity (5.69e-9) and phosphoric ester hydrolase activity (p-value 1.38e-7). In contrast, female GO terms are enriched for peptidase activity (p-value 2.38e-6), serine-type peptidase activity (5.22e-6), serine hydrolase activity (5.22e-6) and amine-lyase activity (8.87e-5). This indicates that females and males express different enzymes at higher levels in adulthood. In a separate analysis, we analyzed the genes from Figure 3B with highest expression in male and highest expression in female (Supplemental Figure 3A). GO terms for highly expressed genes in females and males confirmed that females’ enzymatic activity is related to hydrolase, peptidase and serine activity while males are enriched for kinase activity, and phosphorous processes (Supplemental Figure 3B).

We then used a Principal Component Analysis (PCA) to assess how gene expression of embryos, late post-developmental stages and adults varies across stages (Figure 4C). Principal component 1 (PC1) accounts for 19.7% of the variance across stages and PC2 accounts for 14.2%. Interestingly, PC3 (10.3%) separated all stages into two clusters. One cluster (blue circle) has all the stages that are developing while the other cluster (pink circle) has all the stages that are fully developed or developmentally arrested (namely infective juveniles (IJs)). GO analysis of PC3 positive values were enriched for catalytic activity, embryo development, testosterone dehydrogenase, female sex differentiation, steroid dehydrogenase, regulation of hormone process and female genitalia development. Meanwhile, PC3 negative values were enriched for kinase activity, ligand-gated ion channel, calcium-release channel activity, nicotinate nucleotide metabolic process, NAD metabolic process and regulation of autophagy (supplemental Figure 4). In summary, gene expression profiling of *S. carpocapsae 11* post-development stages shows that starting at 24 hours nematodes are differentiating their gonads and are transcriptionally different from early post-developmental stages. We also found that a gene expression comparison between female and male reveals catalytic activity as differentially expressed with females enriched for peptidase and serine activity while males are enriched for phosphatases. Lastly, PCA separated embryos, late post-developmental stages and adults into two clusters based on their developmental activity.

We investigated the conservation of *C. elegans* sex determination pathway in *S. carpocapsae*. We performed an orthology analysis of the genes described in Haag 2005 to identify which sex determination genes are conserved in *S. carpocapsae*, including genes with more than one paralog (Supplemental table 2). Out of the 27 *C. elegans* sex determination pathway genes, we found 10 that have one to one orthologs such as *sex-1* and *fox-1*, which are responsible for X dosage counting elements and are female-promoting (Hodgkin et al., 1994). Another 7 genes have one to many paralogs such as the case of *sdc-1*, the *her-1* transcriptional repressor, which have three homologs; however, as expected, *sdc-2* and *sdc-3* do not have orthologs in *S. carpocapsae* (Klein and Meyer 1993; Lieb et al., 1996; Nonet and Meyer 1991). Interestingly, *mog-4* and *mog-5*, which are global repressors of *fem-3*, are orthologous to the same 5 genes in *S. carpocapsae* with the best hit for gene g2524 (Puoti & Kimble, 2000). Genes *fbf-1* and *fbf-2*, which *are* germline translational repressors of *fem-3*, are orthologous to one gene g11152 which suggests that this duplication happened along the *C. elegans* lineage (Zhang et al., 1997). Lastly, *fog-1*, which is a promoter of spermatogenesis, has one orthologous gene, contrary to *fog-3* which has orthologs in multiple nematodes species but not in *S. carpocapsae* (Chen et al., 2000; Jin et al., 2001). In summary, we found 19 out of the 27 genes to *C. elegans* in the sex-determination pathway, which might be the core genes important in the nematode sex determination pathway.

## Discussion

In this study we improved the genome of *S. carpocapsae* ALL strain with PacBio technology and BioNano Iris system and identified chromosome X. The *S. carpocapsae* genome is in 16 scaffolds with an N50 of 7.36Mb. The sum of the 10 largest scaffolds’ lengths achieves ∼90% of the genome size, while the top 4 scaffolds cover 50% of the genome. In addition, we used stage-specific developmental transcriptomes to re-annotate our new assembly and predicted 30,957 genes, which we used to infer patterns of rearrangements between scaffolds based on macrosyntenic analysis compared to *C. elegans*. This also allowed us to map the expansion of metalloproteases clusters and to identify the set of genes differentially expressed between males and females.

*S. carpocapsae* has 4 autosomal and one sex chromosome (Curran 1989). Chromosome-level macrosynteny revealed that *S. carpocapsae* chromosome X is orthologous to both *C. elegans* and *P. pacificus* chromosome II and IV. In addition, *S. carpocapsae* chromosome X has a small 3Mb section orthologous to *P. pacificus* and *C. elegans* chromosome X. The small 3Mb of *S. carpocapsae* chromosome X section is the only section orthologous to *P. pacificus* chromosome X, while most of the scaffolds are orthologous to *P. pacificus* chromosome I. This small segment of chromosome X orthologous among *S. carpocapsae, C. elegans* and *P. pacificus* might represent a Nigon unit, which are proposed to be deeply conserved linkage groups found conserved between nematodes (Tandonnet et al., 2018). A comparison of *A. rhodensis* chromosomes to four *Caenorhabditis* species including *C. elegans* pinpointed these Nigon units, one of which (NX) was found as a sole component of chromosome X or found combined with other Nigon units (Tandonnet et al., 2018). Interestingly, *S. carpocapsae* scaffolds are orthologous to single *C. elegans* chromosomes, which is reminiscent of a macrosynteny analysis in other nematode clades that also found conservation of orthologous chromosomal parts (Fradin et al., 2017). If the last common ancestor among clades had 6 chromosomes, then it is likely that *S. carpocapsae* had a fusion of *C. elegans* chromosomes II and IV, which also occurred in unichromosomal nematode *Diploscapter pachys* in which the order of chromosomes fused were I, X, III, II, IV, V (Fradin et al., 2017). The conservation of synteny of *S. carpocapsae* to both *C. elegans* and *P. pacificus* highlights the large-scale chromosomal rearrangements in nematode genome evolution. High quality genomes of other *Steinernema* species will allow for comparisons of evolution of parasitism in this genus. Thus, the nematode field would continue to benefit from assembling high quality genomes which would further the study of parasitology.

Differential expression analysis between males and females uncovered an interesting divergence in catalytic activity. Females have high levels of hydrolase activity, serine-peptidase activity, and proteolysis. Similar catalytic activity is found in adult parasitic females (PF) of *Strongyloids* species (Hunt et al., 2016). *Strongyloids* species have a free-living generation with gonochoristic females and males, and a parasitic female generation that lives inside the mammalian host (Hunt et al., 2016). Although, in *Steinernema*, there are female sex-bias in emerging IJ populations little is known about the role of Steinernema females in infecting a host populations (Alsaiyah et al., 2009). On the other hand, the catalytic activity for males was enriched for phosphorylation, phosphorous activity and kinase activity. Protein phosphorylation and dephosphorylation regulates protein function with phosphatases important in spermatogenesis and regulation of sperm motility (Cottee et al., 2004; Guillermet-Guibert et al., 2015). Kinases are important in nematode male fertility and a target of drugs to treat human parasites (Guillermet-Guibert et al., 2015; Nolan et al., 2004). The role of females in infecting a host and the molecular biology of reproduction processes in EPNs are largely unexplored. The study of the reproductive and developmental processes in parasitic nematodes will be important because it could lead to better methods for biocontrol and for mammalian parasite control through the disruption or interruption of the reproductive cycle.

In conclusion, we have improved the *S. carpocapsae* genome from 1,578 to 16 scaffolds and identified its chromosome X. A macrosynteny analysis found that chromosome X is orthologous to *C. elegans* and *P. pacificus* chromosomes II and IV. Our results point to a conserved region of chromosome X among the three species. We also found catalytic activity differences between adult females and males. Further analysis will be required to assess the role of female adults in EPN infection and regulatory relationships of male development. Lastly, we believe the improved genome of *S. carpocapsae* will advance the field of comparative nematode genomics and allow for the mining of new insights in the evolution of nematode parasitism.

## Materials and Methods

### Axenic *S. carpocapsae* culture

*S. carpocapsae* IJs were cultured on *Xenorhabdus nematophila* colonization defective mutant (HGB315) bacteria to produce axenic IJs. The IJs were first cultured *in vivo* in *Galleria mellonella* (waxworms) and surface sterilized (Gaugler and Kaya, 1990). *X. nematophila* bacteria (HGB315) were cultured in tryptic soy broth (cat. No. A00169, BD Scientific) and shaken at 220rpm in a 26°C incubator overnight. Cultures were plated on lipid agar (LA) plates (8 g/L of nutrient broth, 5 g/L of yeast extract, 2 g/L of MgCl2, 7 ml/L of corn syrup, 4 ml/L of corn oil, and 15 g/L of Bacto Agar), using 600μL of culture per plate. Approximately 300,000 IJs were plated across 20 lipid agar plates (∼15,000 IJs/plate) seeded with lawns of *Xenorhabdus nematophila*. Axenic IJs were collected from LA plates after 2-3 weeks by placing the LA plates in larger petri plates filled with DI water. The IJs were collected from the DI and surface sterilized (Gaugler and Kaya, 1990). IJs were stored in Ringer’s solution in tissue culture flasks at a density < 15 IJs/μL until enough IJs (∼2 million) were collected for the DNA isolation.

### PacBio DNA isolation

#### Sucrose float to remove dead IJs and bacterial contamination

Axenic IJs stored in the tissue culture flasks were transferred to conical tubes, spun down, and washed 3x with DI water. IJs were re-surface sterilized by adding 7.5 mL of distilled water to the worm pellet and 7.5 mL of egg solution (3 mL DI water, 4.5 mL 1M NaOH, 2.5 mL of fresh Clorox bleach) and incubating for 5 min. The egg solution was immediately removed, and the IJs were rinsed 3 times by centrifuging at 2000 rpm for 1-2 min and then suspended in 7mL of molecular grade water. 7 ml of cold 60% sucrose was added, and the sample was mixed and spun at 50g (630 rpm/660 rpm) for 1min at 4°C, and then immediately at 1150g (3000 rpm /3180 rpm) for 3 min at 4°C. The live IJs were collected from the top layer and transferred to a sterile conical tube and washed 3x (centrifuging at 1650g (3600 rpm / 3810rpm) for 5 min) with molecular grade water.

#### Grinding of the IJs with a mortar and pestle in liquid nitrogen to break nematode cuticles

A mortar, pestle, and four ultracentrifuge tubes were wrapped in aluminum foil and autoclaved for at least 20 min at 121°C. The cleaned IJs were added to the autoclaved mortar and liquid nitrogen was poured into the mortar and ground with the pestle until the liquid nitrogen evaporated. IJs were ground for 10 min adding liquid nitrogen as needed. The ground powder was transferred into a 50mL conical tube on ice.

#### IJ cell/tissue lysis and protein and RNA digestion

In a 15 mL conical, 9.5 mL Qiagen Buffer G2 from the Genomic DNA Buffer Set (Cat. No. 19060, Qiagen), 19 μL RNase A (100mg/mL) (Cat. No. EN05231, Thermo Fisher) and 375 μL of proteinase K (> 800mAu, Cat. No. P4850-5ML, Sigma-Aldrich) were mixed, added to the ground IJ sample, and incubated at 50°C in a water bath for 3-4 hours until the lysate was clear. After incubated the sample was spun down at 5,000xg for 10 min at 4°C to separate out any particulate matter that can clog the genomic tip column. The supernatant was transferred to a clean 15 mL conical tube.

#### Genomic DNA measurements

The total amount of genomic DNA was measured with the Qubit fluorometer to determine which genomic tip size to use. We used the 100G genomic-tip because we had between 80-100 μg of gDNA.

#### Genomic-tip protoco

A 100G Qiagen Genomic-tip (100/G) was equilibrated with 4 mL of Buffer QBT. Separately, the DNA sample was diluted with an equal volume of buffer QBT and vortexed for 10s at maximum speed and immediately applied to the equilibrated genomic-tip. The following steps of the genomic-tip were followed according to the manufacturer’s instructions, except that the ethanol and isopropanol precipitation centrifugation steps were performed at 10,000xg for 45 min each. The DNA was eluted in EB buffer overnight at 55°C. The DNA was sheared using 10 pumps of a blunt 24-gauge needle followed by 10 pumps using a blunt 21-gauge needle. The gDNA concentration, absorbance ratios, and fragment size range were determined before SMRT-bell library preparation.

### *S. carpocapsae* genome assembly

The *S. carpocapsae* genome was assembled with Illumina reads and PacBio reads. Then, further improved in contiguity with BioNano genomics Iris system. Previously published Illumina libraries were assembled into high quality contigs using the Platanus (version 1.2.4) command *platanus assemble* with a k-mer size of 51 (-k 51) and a minimum k-mer coverage cut-off of 5 (-c 5) (Dillman et al., 2015; Kajitani et al., 2014). Then, we sequenced six SMRT cells on the PacBio RS II and obtained a total of 500,026 reads (616,611 sub reads) with a N50 read length of 18,308 bp and a mean subread length of 13,036 bp. This translated to 76.8X PacBio read coverage of the 85 Mb genome. All PacBio reads (76.8X genome coverage) were used with the PBcR pipeline into a self-assembly (PB assembly) (Chakraborty et al., 2015; Koren et al., 2012). Then, a hybrid assembly was assembled using the longest error-corrected PacBio reads that provided 25X coverage of the genome, which was output from the PBcR pipeline, with the Illumina assembly. The hybrid assembly was generated using the following DBG2OLC parameters: k 17, KmerCovTh 10, MinOverlap 100, AdaptiveTh 0.02, PathCovTh 3, RemoveChimera 1. The hybrid assembly and self-assembly was corrected for errors and chimeras using the Illumina and PacBio reads with quiver from the smrtanalysis package from PacBio (version 2.3.0p5) and subsequently Pilon (version 1.16) (Walker et al., 2014). Lastly, the PB-assembly and hybrid assembly were merged with quickmerge and MUMmer (version 3.23) using the default settings to generate a *merged assembly* (Chakraborty et al., 2015; Kurtz et al., 2004).

#### BioNano

DNA was extracted from *S. carpocapsae* Infective Juveniles (IJs) using the Animal Tissue DNA Isolation kit (Bionano Genomics). Bionano Irys optical data was generated and assembled with IrysSolve 2.1. We then merged the Bionano assembly with the merged assembly using IrysSolve, retaining Bionano assembly features when the two assemblies disagreed (Supplemental Figure 5A).

The new *S. carpocapsae* genome assembly also went through Haplomerger to create an assembly with minimum possible number of haplotypes. First, the assembly was soft-masked with Windowmasker (version 2.2.22) (Morgulis et al., 2006). Next, the soft masked genome was cleaned using the faDnaPolishing.pl script provided by HaploMerger2 (Huang et al., 2012). Then, the longest 5% of the genome was used as a target and the remaining 95% was used as a query for alignment with Lastz, which created a score matrix. The alignment threshold was kept at 95% to best identify heterozygosity. Lastly, *S. carpocapsae* ALL strain was compared to Breton strain through genomeevolution.org using synmap function with default settings (Lyons & Freeling, 2008).

This Whole Genome Shotgun project has been deposited at DDBJ/ENA/GenBank under the accession AZBU00000000. The version described in this paper is version AZBU02000000.

### *de novo* developmental transcriptome assembly with Oases

Smartseq2 RNA-seq datasets spanning 16 developmental stages (zygote, 2-cell, 4-cell, 8-cell, 24-44-cell, 64-78-cell, comma, 1.5-fold, 2-fold, moving, L1, nonactivated IJ, 9-15h activated IJ, L4, male, and female) in replicates of 2-4 were assembled into stage-specific transcriptomes using Oases 0.2.8 and Velvet 1.2.10. Stage-specific transcriptomes were assembled using different kmer sizes from 22-35 in steps of 2, and then merged into a final assembly. The merged final transcriptome assemblies were compiled into a single fasta file to use for gene model prediction.

### Genome annotation training and prediction with Augustus

539 *S. carpocapsae* genes sequences that matched 458 *C. elegans* CEGMA genes (OrthoMCL) were used together with *de novo* assembled stage-specific transcriptomes to automatically train Augustus gene model exon and intron prediction (Stanke et al., 2008). *C. elegans* UTR models provided by Augustus were used for predicting the *S. carpocapsae* UTRs. A compiled set of *de novo* assembled stage-specific transcriptomes (from above) was mapped onto the merged genome using Blat version 36 with the following settings: −maxIntron=70000 –minScore=100 −minIdentity=94. The best alignments were taken using the command pslReps with the setting: −singleHit, and was then sorted and converted into hints file for Augustus using blat2hints.pl from Augustus (version 3.2.1).

### Assessing genome assembly completeness with Benchmarking Universal Single-Copy Orthologs (BUSCO)

Genome completeness was checked with BUSCO v3 software with default settings for genome and using near-universal single-copy orthologs selected from OrthoDB v9 nematoda_odb9 (Simão et al., 2015). Nematode_odb9 has 982 groups for which 854 groups were found in the *S. carpocapsae* assembly, which is an 87% completeness (Supplemental Figure 5B)

### *S. carpocapsae* female and male DNA collection

IJs were cultured with *X. nematophila* for approximately 54 hours to collect 100 males and 62 hours to collect 50 females. Female and male DNA were extracted with DNeasy blood and Tissue kit (Qiagen Cat No. 69504). The DNA of females and males were processed following the steps F through J from protocol of Serra et al., 2018. Briefly, DNA was tagmented using the Nextera DNA library prep kit (Illumina, FC-121-1030). Then, tagmented DNA was amplified using Phusion High Fidelity PCR master mix with the amplification program set to 1) 72°C 5 min. 2) 98°C 30 sec. 3) 98°C 10 sec, 63°C 30 sec., 72°C 1 min. (repeat 10x). 4) 4°C Hold. PCR samples were cleaned with Ampure XP beads with a 1:1 ratio, concentrations measured with Qubit fluorometer and bioAnalyzed with Agilent 2100 Bioanalyzer. Libraries were prepared and sequenced as paired-end, 43 base pair reads on the Illumina Nextseq 500.

### Genome analysis

To calculate coverage of genome for male and female, first, *S. carpocapsae* genome was indexed with bowtie (version 1.0.0) and DNA reads were mapped to the genome with the following options: −X 1500 −a −v 2 --best --strata –S (Langmead et al., 2009). Bam files were used to calculate coverage using DeepTools2 (version 3.1.1) with default settings for 100kb (Ramírez et al., 2016). Next, gene density was calculated and graphed with GenomicFeatures (version 3.8) and KaryoploteR (version 3.8) packages for R/ Bioconductor (Gel & Serra, 2017; Lawrence et al., 2013). GC content was calculated with perl script GC-content-in-sliding-window using default settings for 100kb window (Richard, 2018). Number of repeats were calculated with RepeatModeler (version 1.0.8) and RepeatMasker (version 4.0.7) using default settings (Smit, Hubley, & Green, n.d.).

### Orthology analysis

Orthologs were determined across *S. carpocapsae* and *C. elegans*, and *S. carpocapsae* and *Pristionchus Pacificus* by blasting the longest protein sequence for each gene using OrthoMCL (version 1.4) with the default settings (L. Li, Stoeckert, & Roos, 2003). Protein sequences were downloaded from WormBase Parasite for *C. elegans* (version PRJNA13758) and *P. pacificus* (version PRJNA12644).

### Protease identification, location, and phylogeny

Protein sequences for the new *S. carpocapsae* genome were aligned to the MEROPS (Rawlings et al. 2018) database using command-line BLAST (Camacho et al. 2009). An additional analysis was done by Neil Rawlings using a custom pipeline to search our genome against the MEROPS collection (Rawlings et al. 2018). Matches with e-value greater than 1xe-10 were kept. Locations for serine and metalloprotease genes were obtained from GTF files. Plots for gene locations and clusters were done using ggplot2. Sequences for orthologs in *C. elegans* were obtained as mentioned above to generate phylogenetic trees for serine and metalloproteases. Genes were aligned in MEGA7 using MUSCLE and a maximum likelihood tree was done using bootstrap support of 100 replicates.

### RNA-sequencing of *S. carpocapsae* time-course, male, and female

To collect RNA for the time-course, *S. carpocapsae* IJs (∼10,000) were cultured with *X. nematophila* in a lipid plate for each allotted time: 3, 6, 9, 12, 15, 24, 36, 48 hours, female and male. The nematodes were washed from the plate into a 1.5mL eppendorf tube and subsequently washed 2 times with Ringer’s and 3 times with HyPure water. Then, *S. carpocapsae* single nematode transcriptome sequencing was performed according to Serra. et al., 2018. Briefly, a single nematode was picked, transferred to a PCR tube and cut with a 25-gauge needle. Then, 2 µl of lysis buffer (for recipe see Serra et al., 2018) which has RNase inhibitor and proteinase K was added to the tube and incubated at 65 oC for 10min, 85 oC for 1min. Immediately after, 1 µl oligo-dT VN primer and 1 µl dNTP was added to the tube and incubated at 72 OC for 3min. After the incubation period samples were reverse transcribed and PCR amplified according to Serra et. al. After PCR, samples were cleaned with Ampure XP beads, concentrations measured with Qubit fluorometer and bioanalyzed with Agilent 2100 Bioanalyzer to check for cDNA quality. If samples had a good BioAnalyzer profile, they were tagmented with Nextera DNA Library Prep Kit, amplified with adapters and sequenced.

### RNA-seq analysis of time course, embryos, female and male

*S. carpocapsae* genome was indexed with GTF file with RSEM (version 1.2.25) using the command rsem-prepare-reference(B. Li & Dewey, 2011). Embryo data were downloaded from NCBI GEO datasets according to Macchietto et. al. 2017 (GSE86381). Unstranded, paired-end 43 bp RNA-seq reads for each worm were mapped using bowtie (version 1.0.0) with the following options: bowtie −X 1500 −a −m 200 −S −p 8 --seedlen 25 −n 2 −v 3 (Langmead et al., 2009). Gene expression was quantified with RSEM (version 1.2.25) with the command rsem-calculate-expression. For all analyses, gene expression was reported in Transcripts Per Million (TPM).

The TPM generated by rsem-calculate-expression for *S. carpocapsae* samples were normalized according to groups using the R package limma because samples were collected, processed and sequenced in different batches (Ritchie et al., 2015). For all RNA-seq analysis genes were kept if reached a minimum of 1TPM for three replicates. For heatmap, gene expression was mean-centered, normalized, and hierarchically clustered with Cluster 3.0 and visualized using Java Treeview (de Hoon et al., 2004; Saldanha, 2004). Differential gene expression was determined using edgeR (Robinson et al., 2010). Genes were considered differentially expressed if the FDR < 0.05 and log2(fold change) > 2. Gene ontology enrichment analyses was calculated using Blast2GO fisher’s exact test and considered statistically significant if FDR < 0.01 (Conesa et al., 2005). List of genes used in Blast2GO were differentially expressed according to edgeR or dynamically expressed according to maSigPro. RNA-seq reads and gene expression tables were submitted to Gene Expression Omnibus (GEO) under the accession number [GSE127823].

## Acknowledgements

The authors thank Neil Rawlings for helping with the MEROPS analysis.

---

**Supplemental table 1:** *S. carpocapsae* scaffold sizes and their respective number of genes

| Scaffolds names | Size (bp) | Number of genes |
| --- | --- | --- |
| Chromosome X | 20,922,283 | 6,776 |
| Scaffold 1 | 10,667,315 | 4,178 |
| Scaffold 2 | 7,849,269 | 2,985 |
| Scaffold 3 | 7,362,381 | 2,988 |
| Scaffold 4 | 6,281,995 | 2,258 |
| Scaffold 5 | 5,918,336 | 2,075 |
| Scaffold 6 | 4,503,743 | 1,813 |
| Scaffold 7 | 4,445,334 | 1,594 |
| Scaffold 8 | 4,020,605 | 1,373 |
| Scaffold 9 | 3,654,988 | 1,471 |
| Scaffold 10 | 3,583,767 | 1,503 |
| Scaffold 11 | 1,806,464 | 724 |
| Scaffold 12 | 1,744,715 | 588 |
| Scaffold 13 | 1,168,849 | 426 |
| Scaffold 14 | 375,282 | 133 |
| Scaffold 15 | 207,283 | 72 |

**Supplemental table 2:**
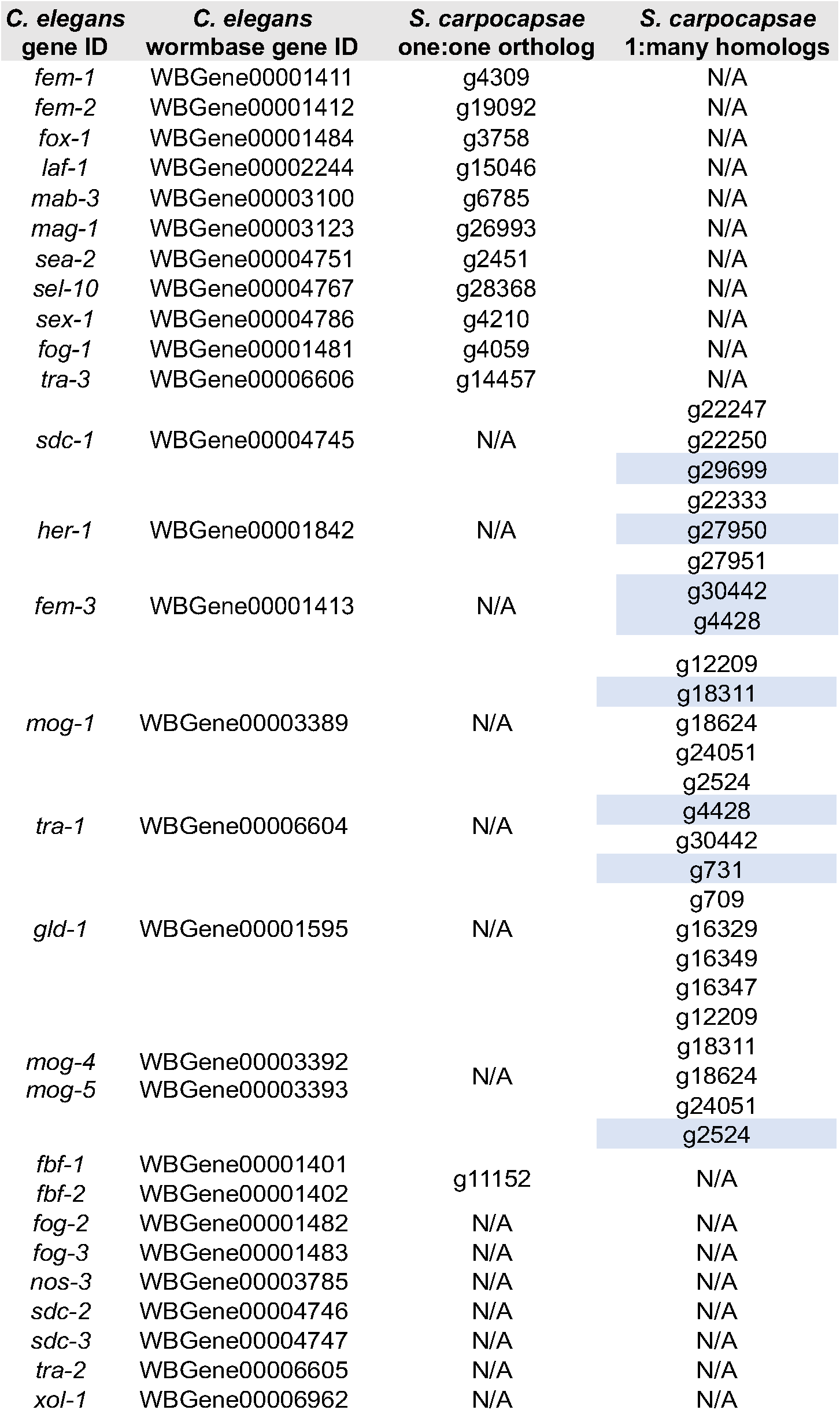
*C. elegans* sex-determination orthologous genes in *S. carpocapsae*. Blue highlighted genes are the best match according to phylogenic analysis.

**Supplemental Figure 2.**
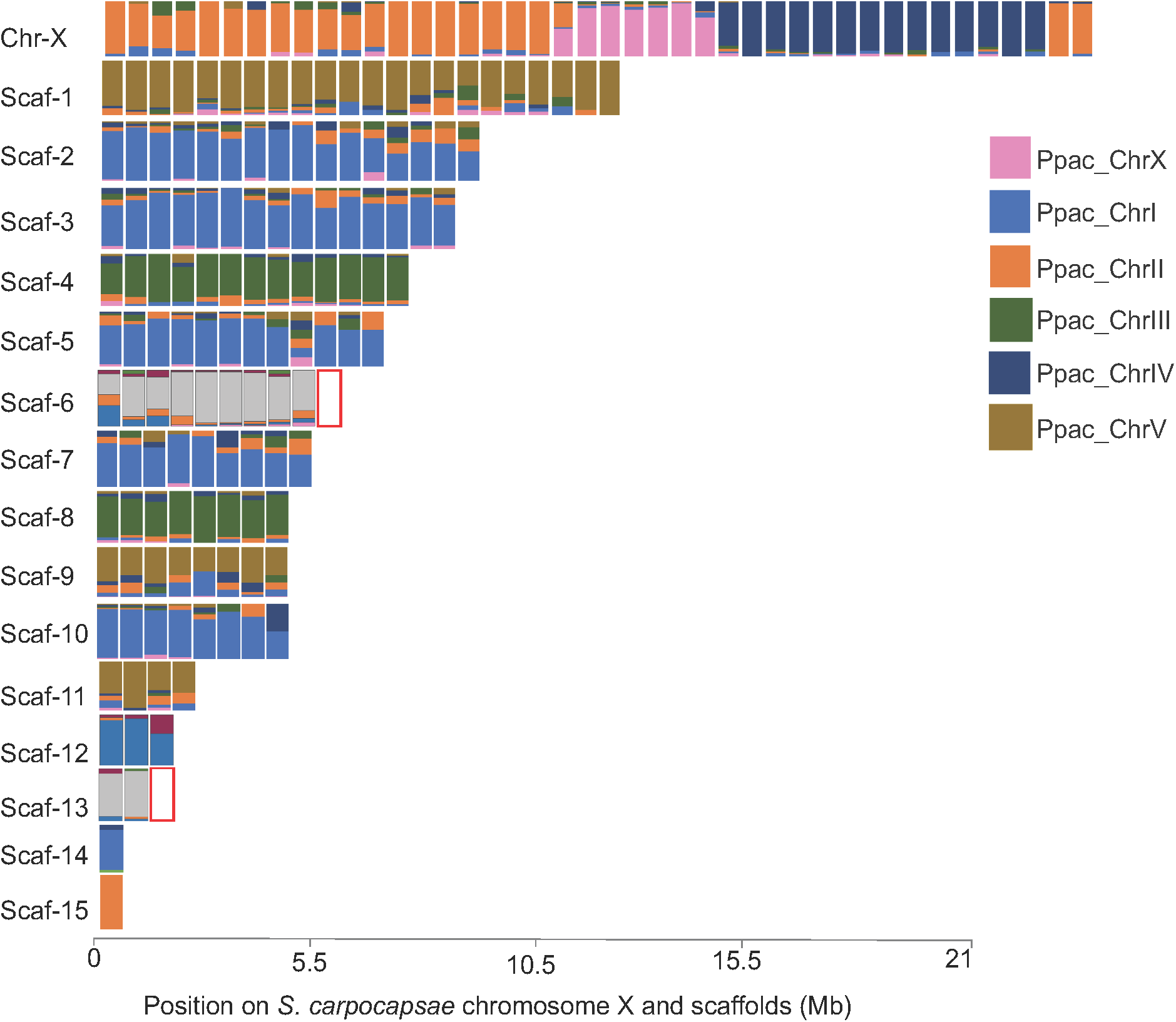
Macrosynteny between *S. carpocapsae* and *P. pacificus*. *P. pacificus?* one-to-one orthologs genes had their position predicted in the *S. carpocapsae* assembly. Each rectangle represents the fraction of *P. pacificus* genes present per 500-Kb window in *S. carpocapsae*. Red rectangle indicates no synteny.

**Supplemental Figure 3.**
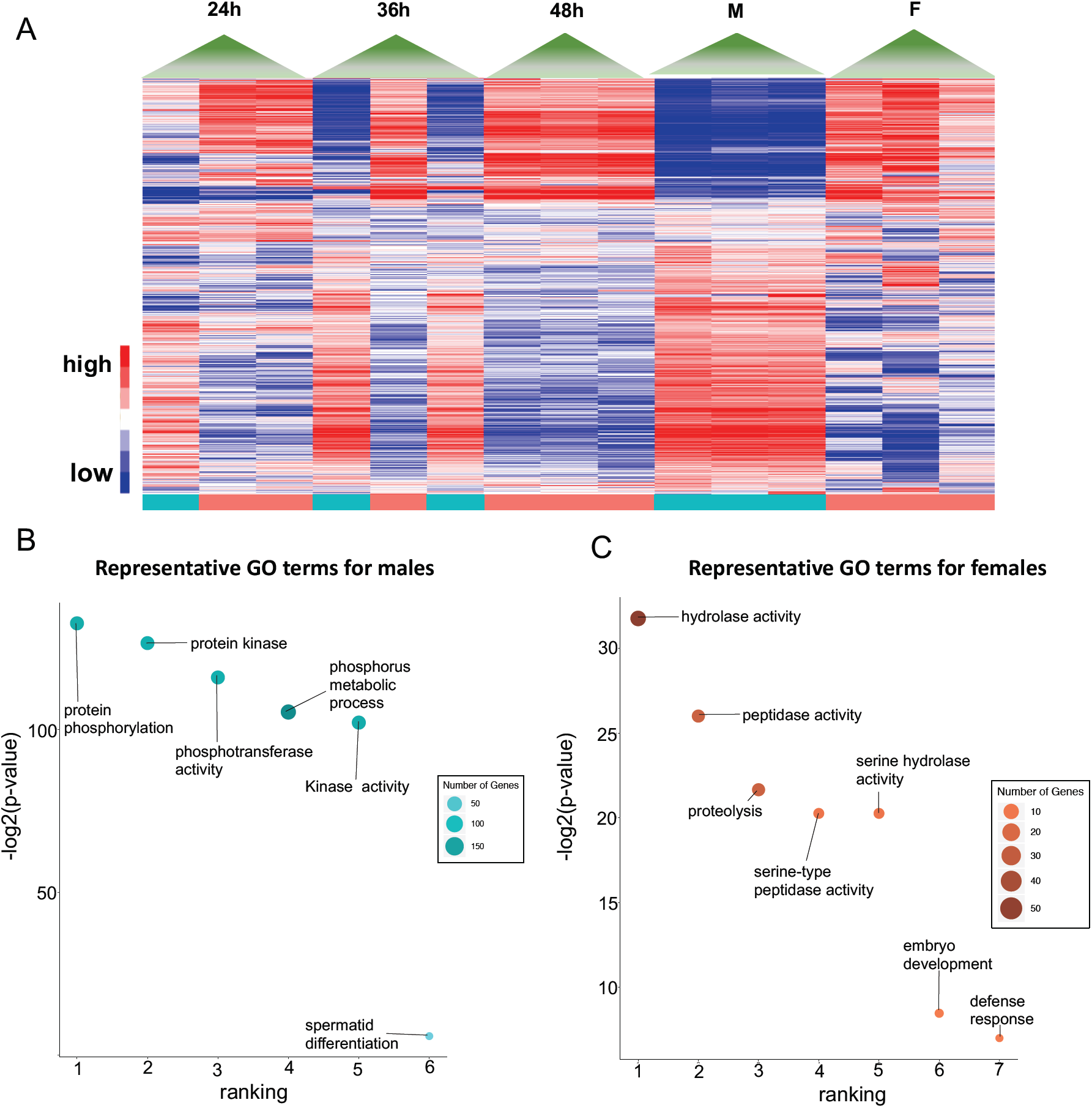
Catalytic activity in female and male adults. **A)** Heatmap of late-stage development and adult males and females for all the genes that are highly expressed in females and males. **B)** Representative GO terms for males from genes highly shown in heatmap. **C)** Representative GO terms for females from genes highly expressed in heatmap.

**Supplemental Figure 4.**
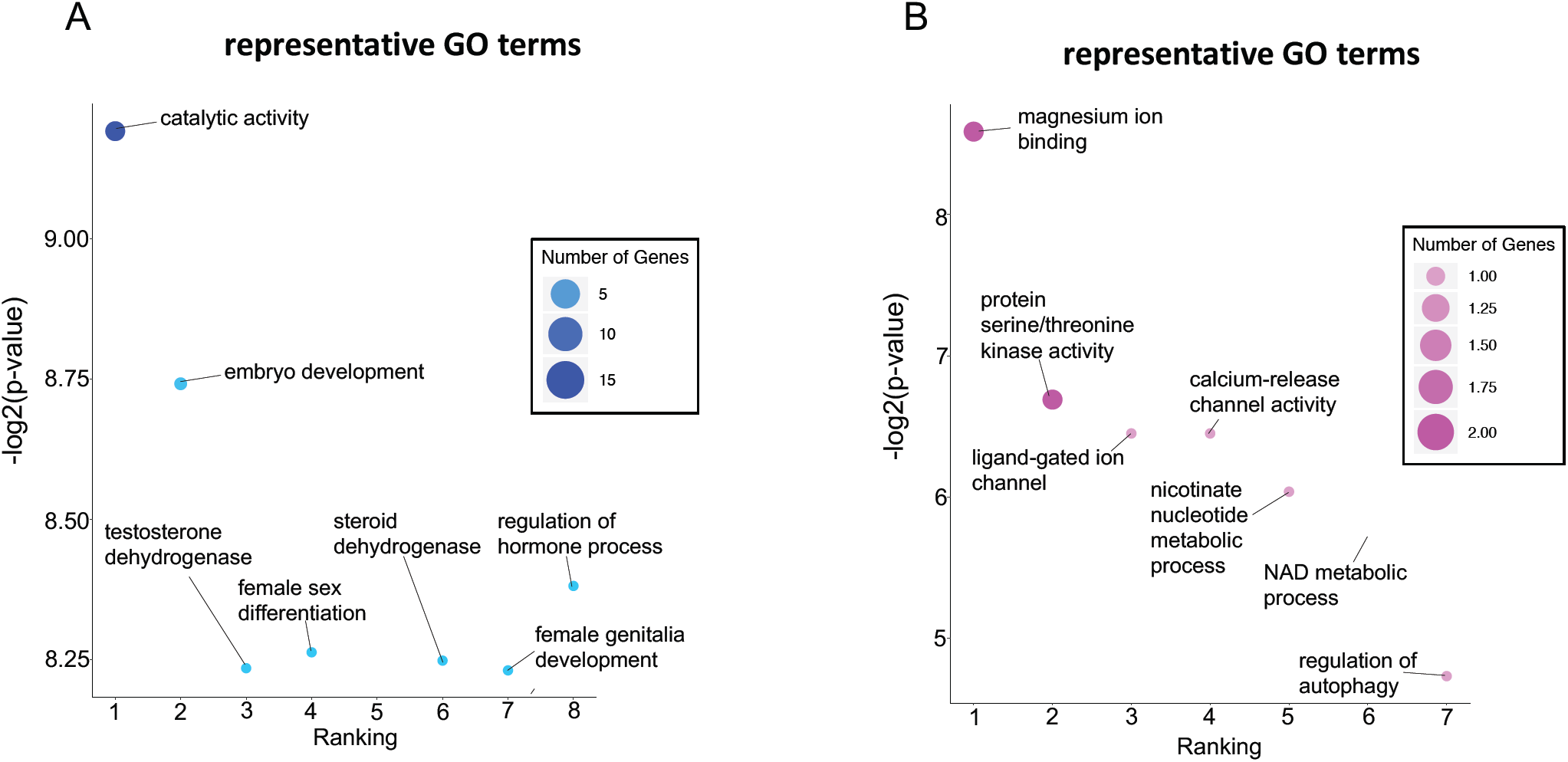
Representative GO terms for PC3 positive and negative rotation values. **A)** Representative GO term for positive value genes (blue). **B)** Representative terms for negative values genes (pink).

**Supplemental Figure 5.**
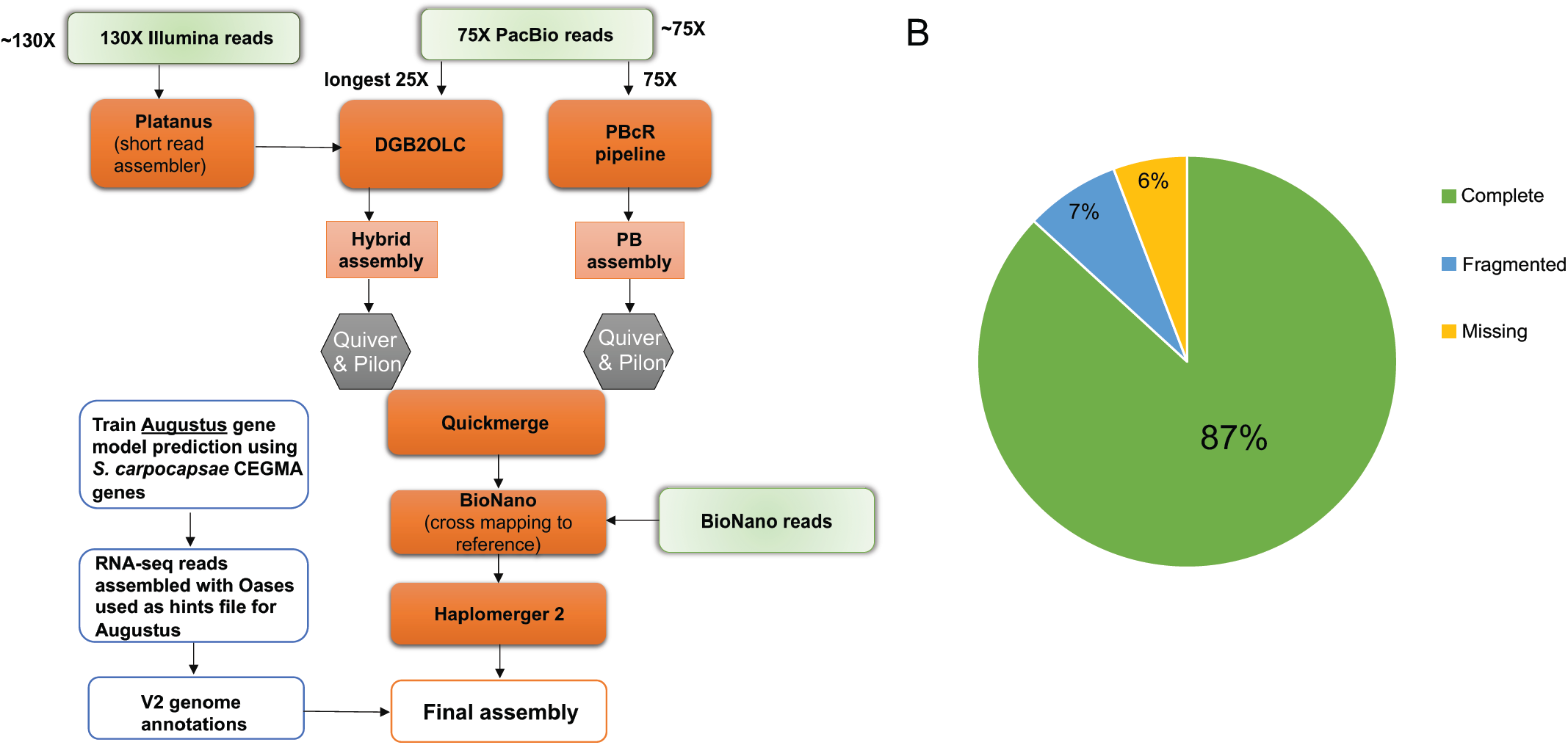
Genome assembly workflow and BUSCO results for *S. carpoc*apsae genome. **A)** An overview of the pipeline used for genome assembly and a brief description of how annotations were generated. **B)** The pie chart illustrates BUSCO results for completed (green), fragmented (blue) and missing (yellow) genes compared to the 982 genes of Nematoda.

